# A novel subtyping method for TNBC with implications for prognosis and therapy

**DOI:** 10.1101/2025.07.04.663242

**Authors:** Zahra Mesrizadeh, Kavitha Mukund, Jovanny Zabaleta, Luis Del Valle, Jerneja Tomsic, Susan Neuhausen, Yuan Chun Ding, Victoria Seewaldt, Augusto Ochoa, Lucio Miele, Shankar Subramaniam

**Author notes:** Corresponding authors: Shankar Subramaniam, PhD; Victoria Seewaldt, MD.

## Abstract

The biological heterogeneity of triple-negative breast cancer (TNBC) poses significant challenges for diagnosis, prognosis, and treatment. While prior TNBC subtype classifications exist, they are not widely used clinically. Here, we aimed to subtype TNBC based on transcriptomic profiles using cell type and state heterogeneity in tumor tissue from 250 pre-treatment women (127 African-American and 123 European-American). We identified three major subtypes and three distinct groups exhibiting unique cell-type composition and mechanisms: Subtype-1 immune signaling/T-cell response; Subtype-2 pro-fibrotic and immune desert; Subtype-3 fatty acid and nuclear receptor signaling. Subtype-1 showed potential responsiveness to immunotherapy, while Subtypes-2 and 3 suggested alternative therapeutic targets. In Subtype-3, which contained a patient group with high *ESR1*, (but not high ERα protein expression) we identified putative mutations in the gene that are unique to these patients. This framework provides a path toward personalized TNBC treatment and is accessible through a user-friendly RShiny application for clinical use.

## Introduction

Triple-negative breast cancer (TNBC) is a biologically heterogeneous and aggressive subtype of breast cancer (BC) that lacks immunohistochemically detectable expression of estrogen receptor (ERα), progesterone receptor (PR), and overexpression/gene-amplification of human epidermal growth factor (*HER2)*^1–4^. TNBC frequently occurs in young parous women who self-reported-race (SRR) as African-American, Black, or African (AA) and in women who harbor a germline *BRCA1* mutation^5^. While much is known about the biology of TNBC, there is a need for a subtyping system that can guide treatment decisions. Recent efforts have focused on classifying tumors based on gene expression profiles and defining a set of gene markers specific for each subtype^6–10^. These subtypes are not typically used to guide treatment-decisions - with the possible exception of anti-androgen therapy^11–14^. Importantly, prior subtyping approaches fail to adequately capture the complex functional and regulatory networks involving interactions between the malignant mammary epithelial cells and the associated TME. Here, we focus our efforts on evaluating the impact of cellular heterogeneity on TNBC biology considering the interactions between the tumor and TME, to obtain unique TNBC subtypes. Our aim is to improve precision therapy, and subsequently, the survival of women with TNBC.

In the companion paper (Hossain et al.), we show that within a diverse TNBC samples (with nearly equal proportions of women who self-reported as AA or EA), the classification scheme developed by Lehmann *et al*.^8^ leads to substantial subtype heterogeneity. Further, Lehmann *et al*.^8^ and other prior approaches fail to classify a group of TNBCs that demonstrate evidence of transcriptional activation of genes regulated by nuclear receptor signaling (NRS) and cell fate determination pathways. We postulate that tumor tissue heterogeneity manifests through distinct combinations of cell type proportions, which can dictate the state of TNBC. The subtypes identified using this heterogeneity information can better classify TNBC, with implications for understanding disease severity and treatment.

To explore our hypothesis and to develop an extensible TNBC classification, we design a novel cell type-driven subtyping approach using tissue-specific gene expression in pre-treatment biopsies of 250 TNBCs from 127 AA and 123 EA women. We utilize breast tissue cell type-specific markers and clustering approaches to identify TNBC subtypes. Using our approach, we identify nine groups within three subtypes for our TNBC samples. Each subtype shows distinct cell type and cell state heterogeneity, prognosis, and survival and is associated with distinct functional mechanisms alluding to the subtype-specific response to targeted therapies.

## Patients and Methods

### Sample description and RNAseq data processing

As described in the companion manuscript (Hossain et al.), 3188 women with TNBC from the Louisiana Tumor Registry (LTR) were selected, of which, 253 TNBC pre-treatment samples were subject to RNA-sequencing (see Data Supplement, tables S1-S2 from Hossain et al.). We note here that we used 250 (127 AA, and 123 EA) samples amongst 253 samples presented in the companion paper after initial processing. These 250 samples are henceforth referred to as LTR TNBC. 227/250 samples were further subject to diversity analysis (genetic admixture testing) using 163 ancestry-informative single nucleotide polymorphisms, as described in Hossain et al.^15^

The quality of raw sequencing data was evaluated using *fastqc v0.11.9* and summarized for downstream analysis using *multiQC*. RNA read mapping and alignment were performed on human reference genome GRCh38.p13_2021v using *Rsubread v2.10.5*, to produce a count matrix containing 44,715 genes and 250 samples, which were filtered for low-count genes. The resulting data matrix containing 23,264 genes across 250 samples was used for all downstream analysis. Count matrix normalization and differential analysis were performed using *DEseq2 v1.36.0* in R, as outlined in their vignette^16^. Differentially expressed genes (DEGs) for every comparison were called at a Benjamini-Hochberg false adjusted *p* < 0.05.

Z-scores for each gene were computed, and genes with a standard deviation (SD) less than ± 2 were filtered to eliminate genes with low biological significance. A total of 18,308 genes passed our significance threshold, and their normalized expression was used as input for visualizations. Heatmaps were generated using the *pheatmap v1.0.12* library in R. The colors on the heatmap represent the normalized expression count scaled row-wise using the built-in scale function available through *pheatmap*.

### Collective Cell Type Markers and subtype Discovery

Cell type markers from twenty-two cell types expected to play a role in breast cancer tumors and tumor microenvironments were identified in the literature **(Supplementary Table 1)**. Group-specific clusters were detected using a hierarchically clustered tree with the dynamic tree cut algorithm at a cut height of 1.3. *ConsensusClusterPlus*^17^ was additionally employed to identify metagroups in an unsupervised manner, with 1000 iterations and the optimal number of clusters (subtypes) (*k*) at 3, determined using the cumulative distribution function (CDF) curve. To identify gene markers uniquely defining each subtype, we converted the z-scores to percentile values using the *pnorm* function in *stats v4.2.0* library in R, adjusting for an exact mean and SD of each sample. Unique subtype gene markers were defined as genes expressed in the top 20% in at least 50% of the subtype’s population. *VennDiagram v1.7.3* and *ggvenn v.0.1.10* were used for all overlap visualizations.

### Enrichment, visualization, and reference network generation

Functional enrichment was identified via functional annotation clustering available through *clusterProfiler*^18^ or *Enrichr*^19^ in R. Gene ontology (biological process and molecular function) pathway analyses were performed using *clusterProfiler’s EnrichGO* function. A network of very high confidence (>0.85) human PPI was downloaded from STRING database v11.5^20^, containing 12,402 proteins and 139,384 interactions. Additionally, TF-target data downloaded from TRRUST-db^21^ was mapped onto the PPI interaction network to generate a custom PPI/TF network. Subtype-specific PPI/TF-target networks were extracted using subtype-specific DEGs resulting in the CC1-specific network containing 470 proteins with 1,140 interactions, the CC2-specific network containing 276 proteins and 557 interactions, and the CC3-specific network containing 859 proteins and 4,564 interactions. These networks were further customized to capture the functional mechanisms underlying each subtype by selecting all DEGs, then reducing to the TF-targets and cell type markers with their 1-step neighborhood. All networks were constructed, annotated, and analyzed in Cytoscape^22^.

### Survival Analysis

Overall survival (OS) was compared using Kaplan Meier plots and log-rank tests using *survival v3.7-0* and *survminer v0.4.9* in R. OS was defined as the time from diagnosis to death from any cause.

### Variant Calling

Single nucleotide variants (SNVs) and indels were called in all samples from CC1, CC2, and CC3 subtypes, focusing on the ESR1 chromosome locus using *FreeBayes* v1.3.5. Variants were filtered based on quality (QUAL>20) and depth (INFO/DP>100). The filtered variants were annotated using snpXplorer to identify their functional consequences.

### RShiny App

The subtyping approach developed in this study is deployed as an interactive RShiny app for prospective samples and is publicly available via [https://www.tnbcworkbench.org/shiny/TNBCsubtyping].

### Validation Strategy

TCGA-BRCA raw count data for 846 tumors was downloaded using TCGAbiolinks in December 2022 and used for validation of our patient subtyping approach, using the unique markers identified. Samples were characterized into Luminal A (n = 586), Luminal B (n = 107), and TNBC (n = 153) based on their IHC/FISH receptor status. Normalized gene expression data for testing datasets were correlated (Pearson) to the centroids of the unique gene markers for each TNBC subtype identified above. Each testing sample was assigned to the TNBC subtype with the highest correlation.

## Results

### TNBC subtyping based on tissue heterogeneity defined by cell types and states

We obtained RNA sequencing data from pre-treatment breast tissue biopsies of 250 racially diverse women with TNBC from LTR, including 123 EA and 127 AA patients (see Methods). Post-processing was performed as outlined in Methods, and the resulting data containing 23,264 features across 250 samples was utilized for all downstream analysis. For cell type marker extraction, we further reduced the dataset by a z-score transformation (±2 SD) resulting in 18,308 genes across 250 samples (See Methods).

To better understand the subtype distribution in our samples, we first applied the well-established TNBCtype framework developed by Lehmann *et al.*^8^, which classifies TNBC into six distinct subtypes based on specific gene transcripts: basal-like (BL1 and BL2), immunomodulatory (IM), mesenchymal (M), mesenchymal stem-like (MSL), and luminal androgen receptor (LAR) subtypes. We observed that thirty-eight of our LTR TNBCs failed to classify into TNBCtype while the remaining samples showed significant subtype heterogeneity **(Fig. 1A)**. Given the subtype heterogeneity and the racial diversity of the samples, we hypothesized that cell type-based markers could better discern tissue state heterogeneity. To examine the cell type heterogeneity, we first identified 261 marker genes for 22 cell types commonly associated with breast tissue **(Fig. 1B; Supplementary Table 1)**^23–33^. Based on the expression of these markers (see Methods) within the LTR TNBCs, we identified 8 distinct clusters via hierarchical clustering, HC1-HC8, henceforth referred to as groups. We additionally utilized consensus clustering (CC) -an unsupervised clustering approach to identify metagroups. CC identified 3 distinct clusters, CC1-CC3, henceforth referred to as subtypes. CC1 comprised distinctly of groups HC1, a part of HC2, and HC3, CC2 contained HC4, HC5, and HC6, and CC3 contained HC7, HC8, and a part of HC2 (henceforth referred to as HC9) **(Fig. 1C)**. Among the subtypes identified, there was no distinction between AA and EA samples **(Supplementary Fig. 1)**. Utilizing xCell and EPIC^34,35^ we estimated cell type distributions within each subtype **(Supplementary Fig. 2A-B)**. We utilized EcoTyper^36,37^ to characterize cellular states for the cell types captured within each subtype **(Supplementary Fig. 2C)**.

**Fig 1.**
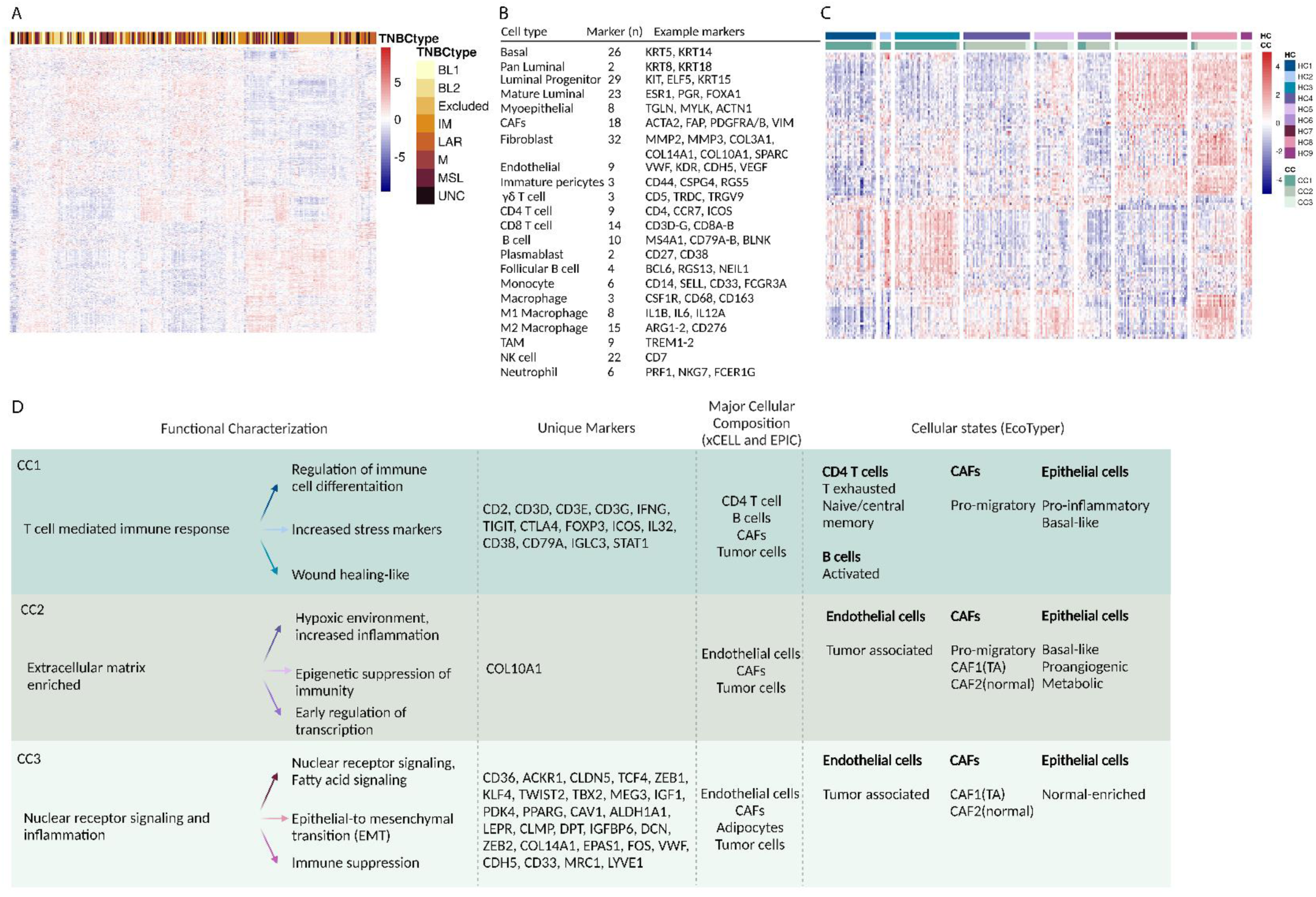
TNBC subtyping and characterization based on cell type marker expression and functional features. (A) The heatmap illustrates the differentially expressed genes for each TNBCtype, as outlined in the Lehmann *et al.*^8^ study. The accompanying bar plot within the heatmap delineates the 7 TNBCtypes, further including the "Lehmann TNBCtype-excluded" group as described in the companion manuscript. Using the Lehmann TNBCtype classification, TNBCs were stratified into 6 subtypes based on significant genes associated with each subtype. (B) The table presents tumor and tumor microenvironment cell types utilized for subtyping LTR TNBCs, along with the number and example of marker genes for each cell type. The complete list of markers is provided in **Data Supplement, Table S1**. (C) The heatmap illustrates relative gene expression (Scaled Normalized Values) of cell type markers within each subtype (ConsensusCluster) and group (HierarchicalCluster) among women with TNBC. This subtyping system categorizes TNBCs into three distinct subtypes and nine groups based on cell type marker expressions. The visualization highlights the heterogeneity between subtypes and within groups, emphasizing the significance of cell type markers. (D) This table encapsulates the major functional features associated with TNBC subtypes and groups identified in the LTR TNBC samples, based on their cell type-associated gene expression. Functional enrichment was performed based on the increased cell type genes in each subtype and group to unveil underlying mechanisms. (From right to left panel: functional mechanism associated with each subtype, functional mechanism associated with each group, unique cell type markers in each subtype, combined results from xCELL and EPIC cellular decomposition, and results from EcoTyper cell state analysis are presented). Arrow colors correspond to the group color scheme from the heatmap in C.

To better characterize the possible functional consequences of the TNBC subtypes, we focused on sets of gene-markers that had increased expression within each subtype and their respective groups (see Methods). In CC1, we identified 42, 65, and 66 gene-markers characterizing HC1, HC2, and HC3, respectively. All three groups within CC1 showed increased expression of transcripts associated with immune signaling and T-cell-mediated response including CD3E-D, CD8A-B, CD7, CD5, GATA3, TCF7, GZMA, PRF1, and PTPRC. Additionally, HC1 showed evidence for T-cell-specific regulation via increased TCF7 and GATA3, while HC2 showed the presence of stress and/or hypoxia markers such as EPAS1 and HIF1A. Lastly, we noted the potential role of wound healing-like mechanism^38,39^ in HC3 as evidenced by increased expression of macrophage markers CD33, CSF1R, TREM1/2, and CD163 as well as extracellular matrix (ECM) markers MMP14, COMP, FN1, PSTN, and COL10A1 **(Fig. 1D)**. Utilizing EcoTyper, we identified that the CC1 subtype is predominantly characterized by infiltrating CD4 T-cells with exhaustion and naïve/central memory state, B cells with activated state, CAFs in a pro-migratory state, and tumor cells in pro-inflammatory and basal-like state **(Fig. 1D)**.

Likewise, in CC2, we identified 27, 68, and 13 gene-markers in HC4, HC5, and HC6 groups, respectively. We observed that all groups in CC2 showed an increased expression of genes associated with ECM including genes such as MMP14, COMP, COL1A1, COL5A1, COL10A1, FAP, and FN1. In addition to ECM markers, HC4 showed increased innate immunity markers including (IL32, TREM1, RUNX1, FCGR3A, MSR1), while HC5 showed the presence of endothelial marker (VWF, CLDN5, CDH5), and HC6 was not associated with a specific functional characteristic **(Fig. 1D)**. The EcoTyper results showed that CC2 had an increased presence of endothelial cells in tumor-associated (TA) states, pro-migratory CAF cells, CAF1 (TA), CAF2 (normal-enriched) state, and tumor cells in basal-like, pro-angiogenic and metabolic state **(Fig. 1D)**.

Similarly, in CC3, we identified 56, 96, and 94 gene-markers in HC7, HC8, and HC9 groups, respectively. We observed that all groups in CC3 were notably enriched for fatty acid (FA) signaling transcripts as evidenced via increased expression of PPARG, CD36, and APOD. Additionally, the HC7 group showed increased gene expression of WNT-signaling-modulator SFRP1, STC2, GATA3, and NRs ESR1 and PGR. HC8 showed increased macrophage genes CD163, TREM1, and MSR1 as well as ECM components COL1A1, COL3A1, COL5A2 and FN1. In HC9 we also observed increased T-cell marker genes CD3E-D and CD8A-B. Consistently, CC3 showed a predominance of adipocytes, pre-adipocytes, endothelial cells with TA state, CAFs with CAF1 (TA) and CAF2 (normal-enriched) states, and tumor cells in a normal-enriched state **(Fig. 1D)**. Our findings suggest that TNBCs can be divided into distinct subtypes with defined functional mechanisms independent of SRR, based on tissue heterogeneity. To ascertain if there exist markers that uniquely associate with SRR we further performed a random forest (RF) classifier with 1000 permutations based on SRR-inferred DEGs. The results were not statistically significant when applied to TCGA TNBC dataset suggesting that the gene markers were sample-specific.

### Functional networks associated with TNBC subtypes

To expand on functional characterizations identified in the earlier section we compared each subtype to all other subtypes (p-adj ≤ 0.05) **(Fig. 2A and Supplementary Fig. 3)** (see Methods). We assessed the functional relevance of each subtype by utilizing custom PPI/TF-target networks (see Methods). Networks constructed from DEGs in CC1 compared to CC2 and CC3 showed immune enrichment including T-cell activation, interferon, and interleukin family members via upregulation of TBX21, STAT1, and IRF1 TFs consistent with the functional characterization of CC1 as noted in the earlier section. TBX21 is associated with CD8 T-cell activation (CD8A, CXCL10, GZMB, LCK, IL12A-B), STAT1 is associated with interferon signaling (IRF1, IRF4, ISG20, OAS2, OASL), and IFNG was associated with upregulation of cytotoxic T lymphocyte markers (CD8A, GZMB) **(Fig. 2B)**. Likewise, DEGs in CC2 compared to CC1 and CC3 showed enrichment for ECM organization pathway genes COL2A1, COL9A1, COL10A1, COL11A1, COMP, MATN3, and MMP13 consistent with the functional characterization of CC2 presented in the earlier section **(Fig. 2C)**. CC3 compared to CC1 and CC2 consistently showed enrichment for FA metabolism and NRS via upregulation of PPARG, ADIPOQ, LEP, FABP4, CD36, and ESR1. (**Fig. 2D**). Also, upregulation of pro-inflammatory genes such as IL6, CXCLs, and SOCS2-3 was associated with FA signaling **(Fig. 2D)**.

**Fig 2.**
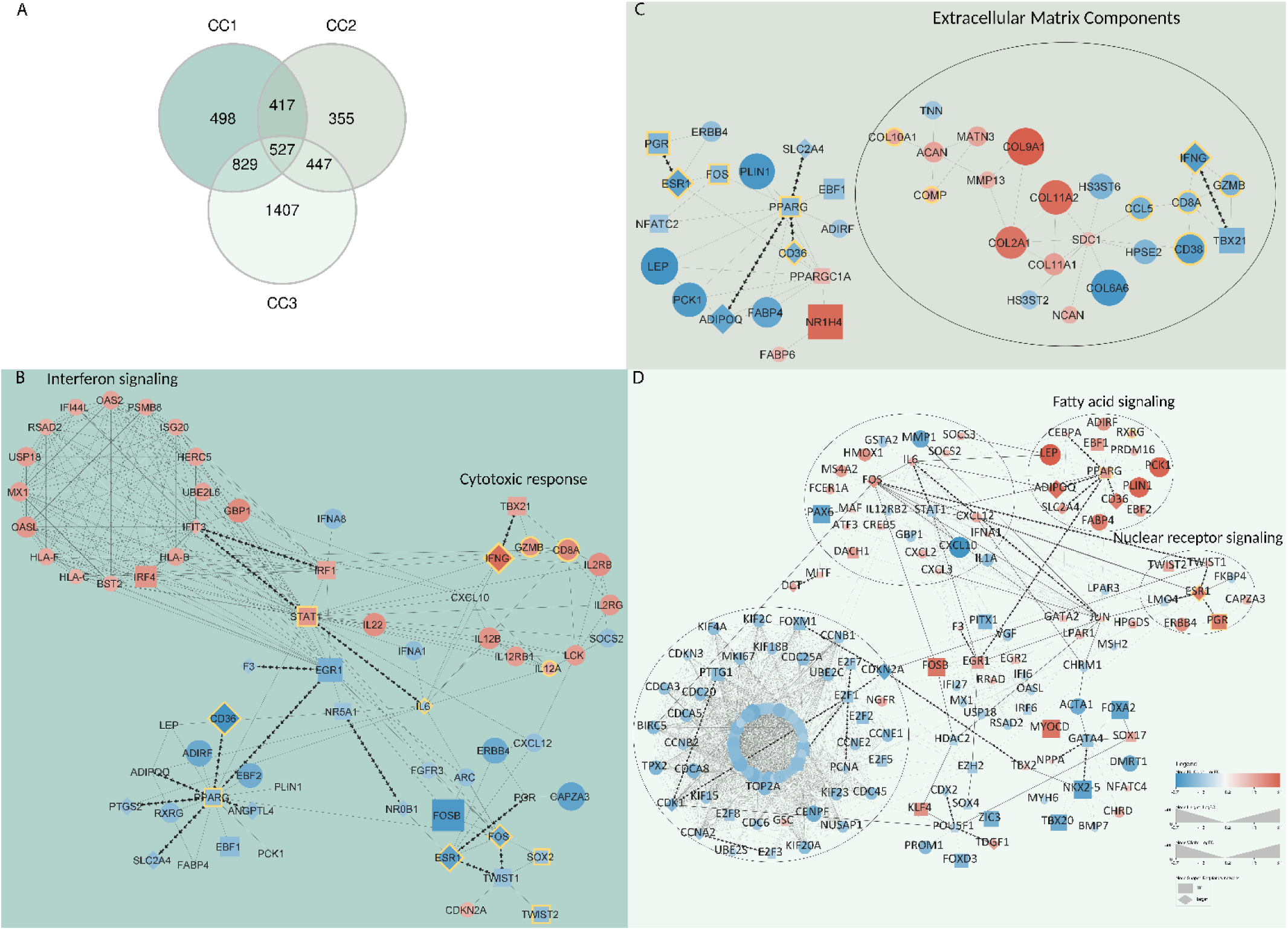
Differentially expressed genes and subtype-specific networks in LTR TNBC. (A) A Venn Diagram illustrates the differentially expressed genes in each subtype. Nodes of each subnetwork are grouped together based on biological mechanism represented in the network, with size and color indicating the log2FC value between subtypes: dark blue (log2FC of -2) to white (0) to dark red (2). (B) A CC1 specific network highlights immune response pathways. (C) A CC2 specific network highlights the significance of extracellular matrix. (D) A CC3 specific network highlights the role of fatty acid, pro-inflammation, and nuclear receptor signaling pathways. The edges present protein-protein interactions, with arrows indicating transcription to target relationships. Yellow border color indicates cell type markers from Fig. 1B.

To expand on ancestry-inferred functional differences in samples with >80% predominant ancestry, we performed DEG analysis comparing AA to EA within each subtype (p-adj ≤ 0.05). In CC1, AA samples showed enrichment for genes involved in immunoglobulin production (IGLV10-54, IGKV1-6) and antigen recognition (TRDV1, TRAV19), while EA showed enrichment for genes involved in stemness and development (LAMA1, FZD5, HOXA4, HOXA9, AXIN2) **(Supplementary Fig. 4A-B)**. In CC2, AA samples showed enrichment for genes involved in humoral immunity (IGHV2-26, IGHV3-48), while EA samples showed enrichment for genes involved in epithelial lineage (KRT1, KRT2, EGFR) **(Supplementary Fig. 4C-D)**. In CC3, AA samples showed enrichment for genes involved in immunoglobulin production (IGLV3-37, IGHD, IGHG4) and antigen presentation via MHC-II (HLA-DOA), while EA showed enrichment for genes involved in epithelial cell processes (SPINK14, KRT85, KRT79) **(Supplementary Fig. 4E-F)**. This is consistent with xCELL data shown in the companion paper, where B-cell lineage enrichment was observed in tumors from AA women as a group, compared to tumors from EA women as a group.

### Subtyping has implications for therapeutic interventions

We estimated likely immune infiltration propensities in each sample using a previously published spatial-phenotype-classifier that provides gene signatures discriminatory of tumor immunophenotypes (excluded, ignored, or inflamed)^40^. The spatial-phenotype-classifier identified CC1 as inflamed, CC2 as excluded, and CC3 as ignored **(Fig. 3A)** consistent with our results as highlighted in the previous sections **(Fig. 1-2)**. Additionally, we characterized the state of T-cell exhaustion (Tex), as previously described from progenitor exhaustion (Tex^prog^, with more stem-like properties) to terminal exhaustion (Tex^term^, with more cytotoxic properties), within LTR TNBC^41^. CC1 was characterized as Tex^term^ showing increased inhibitory receptors and cytotoxic markers while CC3 was characterized as Tex^prog^ with increased TCF7 and low inhibitory markers **(Fig. 3B-C)**. Overall, these results were consistent with our subtype-based functional characterization in the earlier sections and suggest that TNBCs within each subtype might be sensitive to specific therapeutic strategies. In CC1 with increased Tex markers, IFN response, and T-cell activation **(Fig. 1F, 2A)**, immune-checkpoint inhibitors such as LAG3, TIGIT, and CTLA-4 in combination with anti-PD-L1 therapy, may enhance the efficacy of existing chemotherapy regimens^26^. The immune scarcity in the CC2 subtype postulated the potential interventions to prime TNBC for immunotherapies such as anti-TGFB and/or VEGF ^40,43^.

**Fig 3.**
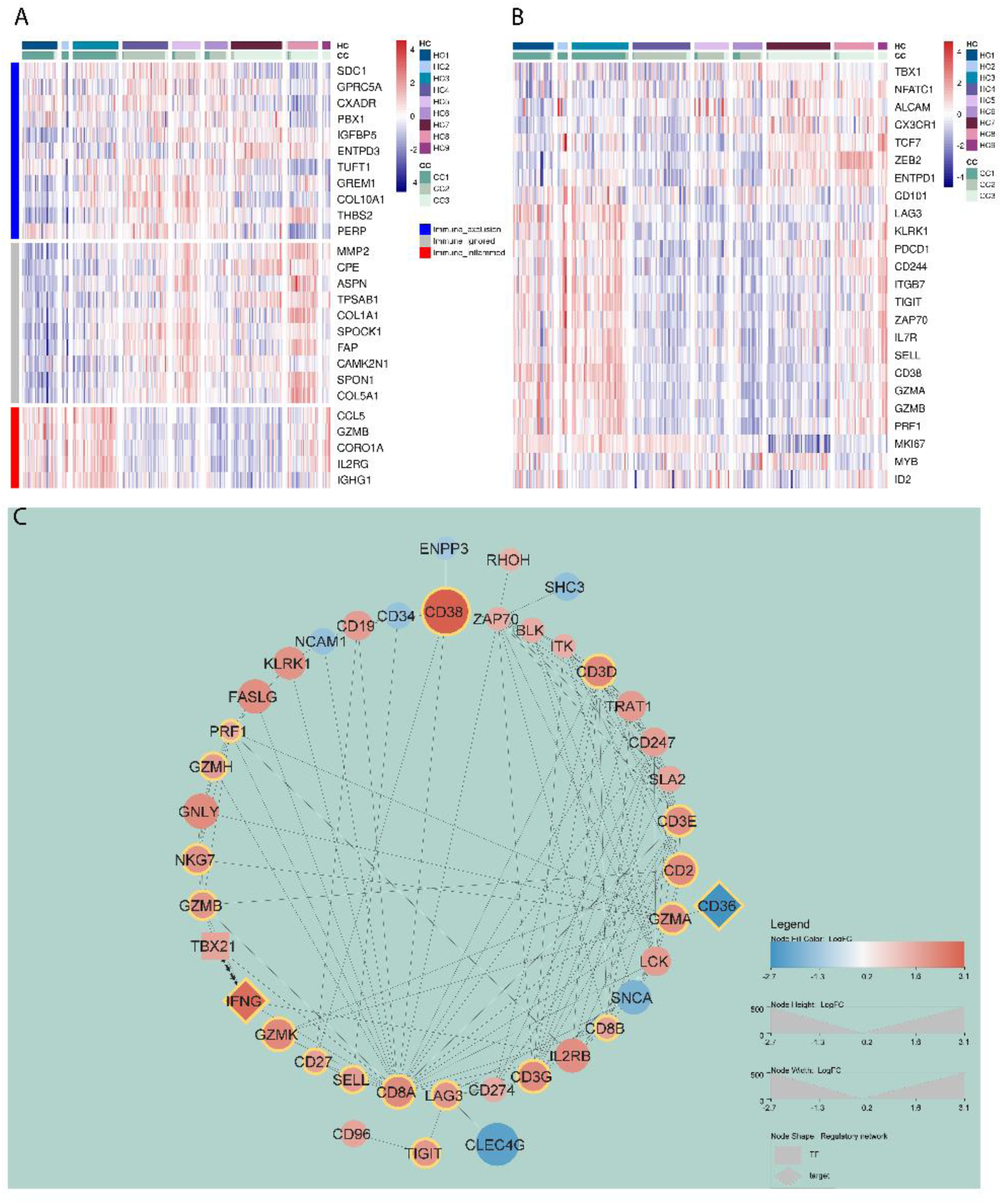
Spatial gene classifiers and T-cell exhaustion across TNBC subtypes. (A) A heatmap illustrates the relative gene expression (Scaled Normalized Values) of spatial-gene-classifier in each subtype (ConsensusCluster) and group (HierarchicalCluster) in the LTR TNBC samples. (B) A heatmap illustrates the relative gene expression (Scaled Normalized Values) of T-cell exhaustion markers in each subtype (ConsensusCluster) and group (HierarchicalCluster) in the LTR TNBCs. (C) A CC1-specific network captures T-cell exhaustion and its neighboring DEGs. Nodes are sized and colored based on the log2FC value between subtypes: dark blue (log2FC of - 2) to white (0) to dark red (2). The edges represent protein-protein/TF-target interactions, with arrows indicating transcription to target relationships. The yellow boarder color indicates T-cell exhaustion markers.

Due to the increased expression of nuclear receptor (NR) transcripts in the CC3 subtype and the availability of drugs targeting NRs, we assessed the known 48 human NR^44^ expression across our subtypes **(Supplementary Fig. 5A)**. We identified 23 NRs, including the entire NR4A subgroup, which showed increased expression in our CC3 subtype **(Supplementary Fig. 5A)**. Interestingly, in the CC3 network, NR4A1 and NR4A2 were associated with the upregulation of cell-fate commitment gene LMX1A as well as FA signaling **(Supplementary Fig. 5B)**. Additionally, steroid sub-family NRs ESR1, ESR2, AR, PGR, NR3C1, and NR3C2 showed increased transcript expression in the CC3 subtype **(Supplementary Fig. 5A)**. These results underscore the subtype’s potential vulnerability to small molecule inhibitors targeting these NRs, providing a promising avenue for therapeutic development.

### Subtyping reveals an ESR mRNA^hi^ (HC7) TNBC group

As mentioned earlier, all TNBCs analyzed were required to have <1% ER expression by IHC. Despite this, transcript mapping of CC3 showed increased expression of ESR1, ESR2, and PGR mRNA (ESRmRNA^hi^) **(Supplementary Fig. 6 and Fig 2D)**. Within the CC3 subtype, 35 TNBCs fell within the HC7 group, all of which were independently identified in the companion manuscript as the "Lehmann TNBCtype-excluded" group, and shown to have a highly distinct transcriptional profile compared to all TNBC-included tumors. To confirm the transcriptional uniqueness of HC7 TNBCs from luminal BC, we 1) compared ESR1 expression across different subtypes and groups in our samples, 2) assessed expression profiles of ESR1 and ESR2 downstream targets in HC7, and 3) compared common features between HC7 and luminal-A BC from TCGA (henceforth referred to as LumA). We observed a higher expression level of ESR1 in CC3 and particularly in HC7 **(Fig. 4A-B)**, while being lower than the luminal subtype **(Fig. 4C)**. We identified that the ESR1 and ESR2 targets were not exclusively associated with CC3 and/or HC7 which indicates the heterogeneous distribution of ESR1 and ESR2 transcriptional activity across our samples **(Fig. 4D)**. Lastly, we observed that the functional enrichment of unique DEGs identified in HC7 (vs. LumA) highlighted the role of the arachidonic acid (ArA) pathway **(Fig. 4E)**, an essential fatty acid required for eicosanoid production (prostaglandins, leukotrienes) and associated with metastasis and pro-inflammation in various cancers including TNBC^45,46^. Collectively, these findings suggest that CC3 represents a TNBC subtype characterized by increased ESR1 mRNA - but not protein-expression with connection to inflammation and FA/eicosanoid signaling based on the constructed network in **Fig. 2D**.

**Fig 4.**
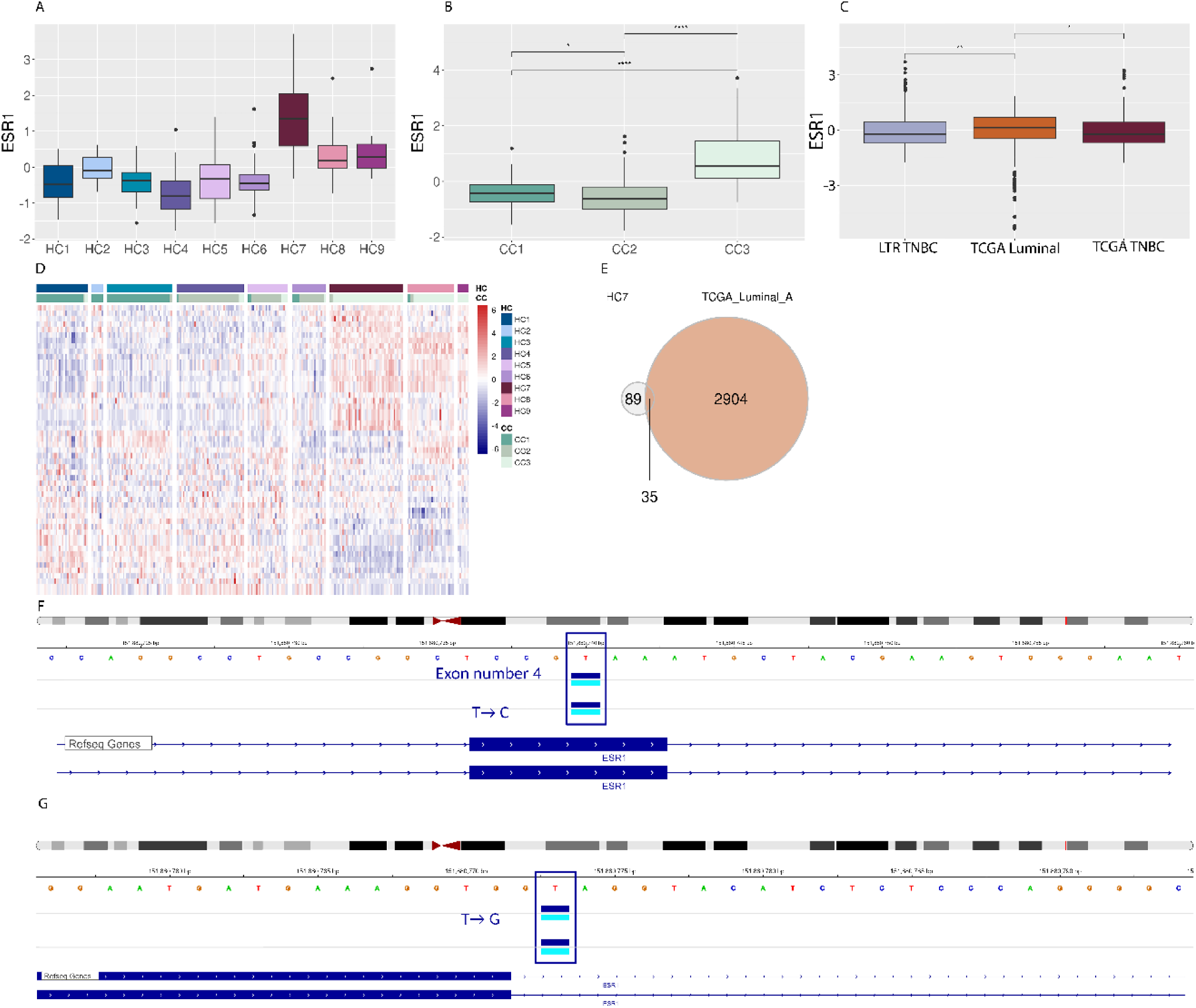
Analysis of ESR1 and ESR2 expression and downstream targets across TNBC subtypes. (A) and (B) Boxplots show the scaled expression of ESR1 and ESR2 in groups and subtypes in the LTR TNBCs. (C) A boxplot shows the scaled expression of ESR1 in the LTR TNBCs compared to luminal and TNBCs in the TCGA dataset. (D) A heatmap displays the relative gene expression of downstream targets of ESR1 and ESR2 in each subtype (ConsensusCluster) and group (HierarchicalCluster) in the LTR TNBCs. (E) Identifying the overlap between genes significantly upregulated in luminal A vs. TNBCs (from TCGA) and genes significantly upregulated in HC7 vs. all others (from LTR TNBCs). (F) and (G) Representative IGV snapshot showing SNVs on chr6 locations in ESRmRNA^hi^ samples (2 examples for each SNV are shown).

Considering the observed upregulation of ESR1 within our samples, we explored if these samples exhibit unique mutations. We identified two single nucleotide variations (SNVs) on chr6 unique to the CC3 subtype: a synonymous codon change at position 151880740 (T-C substitution) and a splice donor SNV at position 151880773 on chr6 (T-G substitution) (see Methods). The synonymous SNV was present in 30/35 ESRmRNA^hi^ samples, and the remaining 5 samples showed a splice donor SNV **(Fig. 4F-G)**. While synonymous SNVs do not result in an amino acid change in the protein they encode, they can affect the corresponding RNA expression level. Splice donors can alter the normal splicing process, potentially leading to the inclusion or exclusion of exons in the mature mRNA influencing protein expression. Further experimental investigations are necessary to understand the role of ESR1 mutations in its mRNA expression in TNBC.

### Subtype-based Survival outcomes and prognostic biomarkers

We examined overall survival (OS) outcomes for each TNBC subtype among patients with stage I-III and younger than 63 years (n_Total_=130, and n_AA_=70 and n_EA_=60). Samples were balanced for stage, age, BMI, and SRR. The analysis revealed that patients classified as CC1 exhibited a favorable prognosis, whereas those classified as CC2 showed notably less favorable prognosis **(Fig. 5A)**. Our analysis identified a set of DEGs with a prognostic value that predicts subtype-specific survival outcomes. Notably, in patients classified as CC1, we found that CD244 is associated with favorable OS, while SLC4A2 is associated with unfavorable OS **(Fig. 5B)**. In CC2, we identified GNLY and IKZF3 as favorable prognostic markers, whereas MIR1262, LOC100419059, GRM5, and TYR are associated with unfavorable OS **(Fig. 5C)**. Especially considering the worse outcomes observed in this subtype, these findings suggest that prognostic biomarkers in CC2 may have potential diagnostic value.

**Fig 5.**
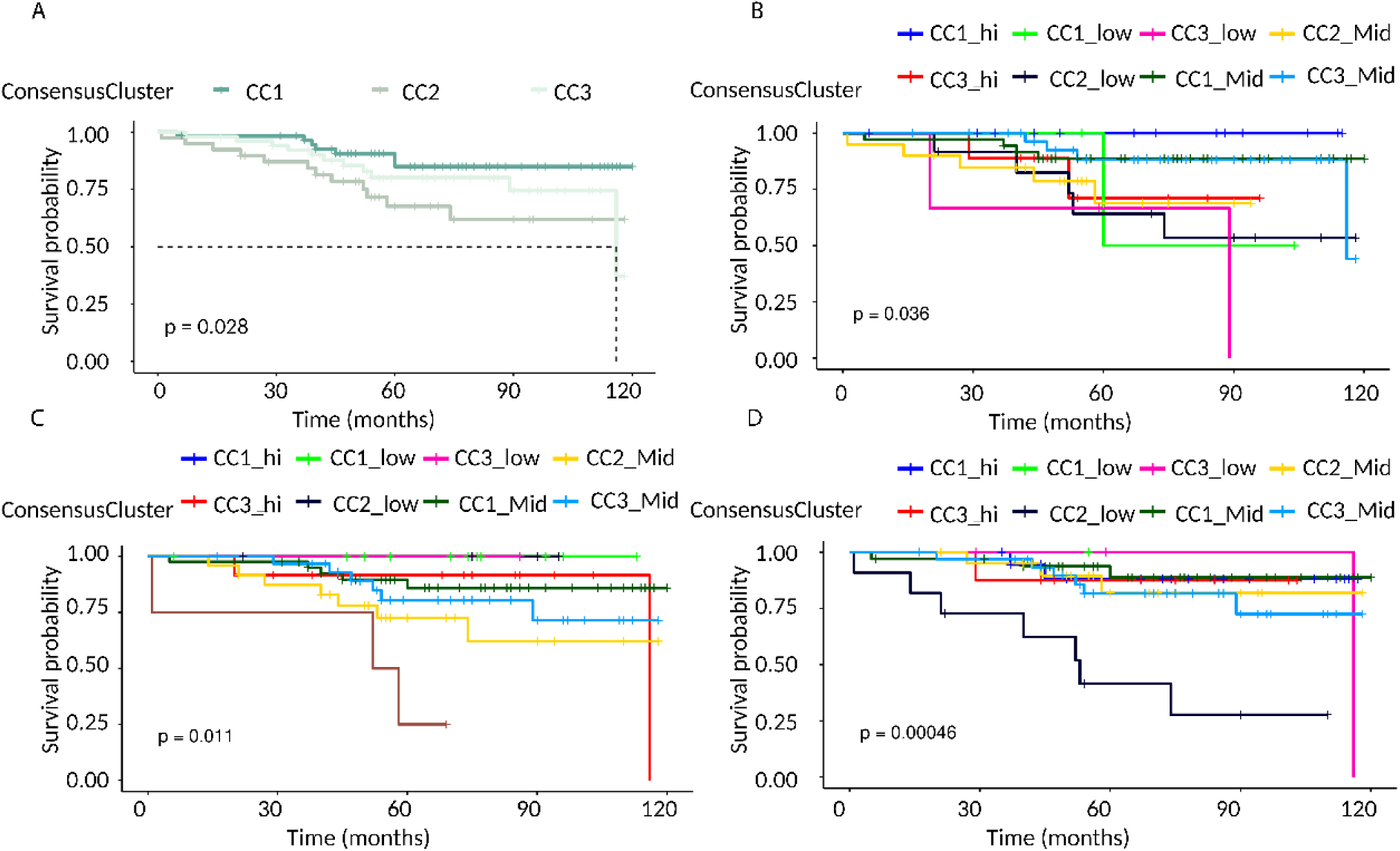
Survival analysis of TNBC subtypes and associated biomarkers. (A) The overall survival associated with each subtype in women under 63 years old with TNBC. (B)-(D) The overall survival associated with CD244, MIR1626, and SLFN12L, respectively, for each subtype in women under 63 years old with TNBC.

Among patients classified as CC3, we identified several genes including PRF1, TNFRSF8, NCR1, IL18RAP, TRGV2, TRGV3, IL2RB, CD79B, CARD11, NCF1, NCF1C, AIM2, FGD2, SIGLEC10, SLC4A2, KCNA5, FBN2, and TYR which are associated with favorable prognosis. In contrast, genes including SLFN12L, GFI1, IKZF3, RNF128, and CRTAM are associated with unfavorable prognosis **(Fig. 5D)**. Interestingly, SLFN12L over-expression in TNBC is associated with sensitivity to chemotherapy^47^ and luminal markers with race-inferred biological differences in TNBC^48^. However, both GFI1^49^ and CRTAM expression have been associated with better prognosis in TNBC indicating further investigation of these biomarkers in the context of CC3 TNBC is necessary to elucidate their role in disease progression.

## Discussion

Classifications of TNBC proposed by Lehmann *et al.*^8^, and Burstein *et al.*^10^, provided valuable insights into TNBC heterogeneity. However, challenges stemming from technical variability, intra-subtype heterogeneity, and issues related to the reproducibility of such classification methods prompted a reevaluation of TNBC subtyping, specifically in the LTR TNBC samples. Our primary hypothesis was that the tumor heterogeneity should be reflected in the tissue and cell heterogeneity and if we can use the latter through cell-specific markers, we will not only be able to capture the tumor heterogeneity but also cellular mechanisms that lend to this heterogeneity. To this end, we developed a novel subtyping approach utilizing 22 breast-specific cell type markers in 250 women with TNBC - with nearly equal distribution of self-reported AA and EA women, who differed significantly for socioeconomic deprivation and obesity rates as described in the companion manuscript. This approach uncovered three subtypes with distinct inferred cell types, cell states, functional-modules, and OS, opening avenues for tailored and targeted therapy. Additionally, ancestry-inferred analysis showed immunoglobulin production plays a dominant role in AA while epithelial state transition mechanisms play a key role in EA TNBC subtypes.

Tumors pose a significant challenge to the immune system, often disrupting the delicate balance between immune activation and suppression, influencing outcomes and prognosis. We identified the CC1 subtype as an immunologically active subtype, characterized by significant immune exhaustion. Furthermore, we discovered CD244, an immunoregulatory receptor expressed on various immune cell types, as a favorable prognostic biomarker. CD244 has been reported to play dual roles in anti-tumor immunity^50^ with evidence for sustaining exhaustion^51^ and as a marker for tumor cytolytic activity^52^-suggesting its complexity as a therapeutic target. The CC2 subtype was significantly associated with the enrichment of ECM components as well as an immunologically “cold” microenvironment. In recent years, studies have highlighted the importance of TME and cell-matrix interactions in understanding tumor development, progression, and metastasis as well as treatment responses^42,53–56^. Hammerl *et al.*^40^ identified COL10A1 as a crucial driver of immune exclusion in tumors (identified as DEG in CC2) alluding to the idea that the observed immune exclusion seen in these patients could be associated with the COL10A1. Studies have demonstrated that ECM-hypoxia crosstalk leads to increased BC aggressiveness^57^, and inflammatory mediators released from TNBC among other factors are involved in upstream regulatory expression of HIF-1α^58^. Consistent with this, we observed an increased expression of hypoxic signatures within the HC4 group within the CC2 subtype compared to all other groups **(Supplementary Fig. 7)**.

The CC3 subtype showed enhanced expression of genes involved in FA metabolism, and NRS **(Fig. 1D, 2D and Supplementary Fig. 3)**. NRs have a well-defined ligand-binding domain and are amenable to small-molecule intervention^59^. NR4A NRs play a diverse role in homeostasis, proliferation, cell migration, apoptosis, metabolite metabolism, DNA repair, glucose utilization, and tumorigenesis^60^. Muscat *et al.* identified NR4A NRs to be associated with both ER+ and ER-BCs compared to normal breast tissue^61^. Additionally, Zhou *et al*., showed that NR4A1 is involved in inducing EMT in a TGF-β-dependent manner, and migration and invasion in an inflammation-dependent manner in breast cancers^62^. Further experiments are necessary to understand the underlying role of NR4A NRs in CC3 TNBC and to utilize these markers to improve patients’ survival outcomes. Furthermore, we observed a set of steroid NRs with increased expression in CC3 **(Supplementary Fig. 5A)**, and as shown in **Fig. 2D**, ESR1 was tightly connected to the upregulation of inflammatory genes as well as FA/eicosanoid signaling. These results alongside ESR1-specific SNVs located on CC3 TNBCs suggest a significant role of NRs in the formation and progression of the CC3 subtype of TNBCs, and further experiments to validate these results are essential. The possible role of ArA metabolites, including eicosanoids, prostaglandins, and leukotrienes, deserves further investigation. PGE2, an abundant ArA metabolite, is one of the most potent immune suppressants in the TME^63^. Tumors of this subgroup may respond to ArA metabolism inhibitors, such as phospholipase A2 (PLA2) inhibitors or cyclo-oxygenase (COX) inhibitors, including NSAIDs. Interestingly, the Miele group and others have reported that NSAID sulindac has activity in immune-competent preclinical models of TNBC, where it enhances tumor immunity^64^. Consistent with these, we observed significantly increased expression of PLA2, COX1, and COX2 within the CC3 subtype **(Supplementary Fig. 8)**.

To enable fellow researchers to utilize our subtyping system, we developed a user-friendly RShiny App, facilitating TNBC subtyping based on tissue heterogeneity and biological mechanisms. This tool helps guide treatment decisions, particularly for immunotherapies like immune checkpoint inhibitors. To ensure the robustness and clinical relevance of our approach, both retrospective and prospective studies are necessary to validate its predictive power for treatment response and survival outcomes in diverse TNBC populations. With further validation, this system could significantly improve personalized treatment strategies for TNBC patients.

## Author Contributions

Z.M. analyzed the data, developed the models, wrote the first version of the manuscript along with preparing Tables and Figures. K.M. provided help with the design of the analysis and building the models. J.Z., L.M. and A.O. were involved in the sample collection with L.D.V. contributing to the clinical assessment and J.Z. contributing the bulk RNA sequencing. All samples were from Louisiana Tumor Registry. S.N., Y.C.D., J.T. and V.S. were involved in the ancestry analysis. S.S. designed and supervised the project and revised the manuscript. All authors have read, edited, and accepted the manuscript.

## Notes

This work was supported by Wellcome Leap Delta Tissue Program (SS, VS), NIH OT2-OD030544, NIH OT2-OD036435, and NIH R01-CA282657 (SS), as well as NCI SPORE P20CA202922 and NIGMS 2U54GM104940 (LM, AO, LDV).

### Competing Interest Statement

The authors have declared no competing interest.

## REFERENCES

1. Derakhshan, F. & Reis-Filho, J. S. Pathogenesis of Triple-Negative Breast Cancer. Annu. Rev. Pathol. Mech. Dis. 17, 181–204 (2022).

2. Garrido-Castro, A. C., Lin, N. U. & Polyak, K. Insights into Molecular Classifications of Triple-Negative Breast Cancer: Improving Patient Selection for Treatment. Cancer Discov. 9, 176–198 (2019).

3. de Ruijter, T. C., Veeck, J., de Hoon, J. P. J., van Engeland, M. & Tjan-Heijnen, V. C. Characteristics of triple-negative breast cancer. J. Cancer Res. Clin. Oncol. 137, 183–192 (2010).

4. Foulkes, W. D., Smith, I. E. & Reis-Filho, J. S. Triple-negative breast cancer. N. Engl. J. Med. 363, 1938–1948 (2010).

5. Prakash, O. et al. Racial disparities in triple negative breast cancer: a review of the role of biologic and non-biologic factors. Front. Public Health 8, 576964 (2020).

6. Martini, R. et al. African Ancestry–Associated Gene Expression Profiles in Triple-Negative Breast Cancer Underlie Altered Tumor Biology and Clinical Outcome in Women of African Descent. Cancer Discov. 12, OF1–OF22 (2022).

7. Roelands, J. et al. Ancestry-associated transcriptomic profiles of breast cancer in patients of African, Arab, and European ancestry. Npj Breast Cancer 7, (2021).

8. Lehmann, B. D. et al. Identification of human triple-negative breast cancer subtypes and preclinical models for selection of targeted therapies. J. Clin. Invest. 121, 2750–2767 (2011).

9. Lehmann, B. D. et al. Refinement of Triple-Negative Breast Cancer Molecular Subtypes: Implications for Neoadjuvant Chemotherapy Selection. PLOS ONE 11, e0157368 (2016).

10. Burstein, M. D. et al. Comprehensive Genomic Analysis Identifies Novel Subtypes and Targets of Triple-Negative Breast Cancer. Clin. Cancer Res. 21, 1688–1698 (2014).

11. Gucalp, A. et al. Phase ii trial of bicalutamide in patients with androgen receptor–positive, estrogen receptor–negative metastatic breast cancer. Clin. Cancer Res. 19, 5505–5512 (2013).

12. Bonnefoi, H. et al. A phase II trial of abiraterone acetate plus prednisone in patients with triple-negative androgen receptor positive locally advanced or metastatic breast cancer (UCBG 12-1). Ann. Oncol. 27, 812–818 (2016).

13. Traina, T. A. et al. Enzalutamide for the treatment of androgen receptor–expressing triple-negative breast cancer. J. Clin. Oncol. 36, 884–890 (2018).

14. Barton, V. N. et al. Multiple molecular subtypes of triple-negative breast cancer critically rely on androgen receptor and respond to enzalutamide in vivo. Mol. Cancer Ther. 14, 769–778 (2015).

15. Pakstis, A. J. et al. SNPs for a universal individual identification panel. Hum. Genet. 127, 315–324 (2009).

16. Love, M. I., Huber, W. & Anders, S. Moderated estimation of fold change and dispersion for RNA-seq data with DESeq2. Genome Biol. 15, 550 (2014).

17. Wilkerson, M. D. & Hayes, D. N. ConsensusClusterPlus: a class discovery tool with confidence assessments and item tracking. Bioinformatics 26, 1572–1573 (2010).

18. Yu, G., Wang, L.-G., Han, Y. & He, Q.-Y. Clusterprofiler: an r package for comparing biological themes among gene clusters. OMICS J. Integr. Biol. 16, 284–287 (2012).

19. Kuleshov, M. V. et al. Enrichr: a comprehensive gene set enrichment analysis web server 2016 update. Nucleic Acids Res. 44, W90–W97 (2016).

20. Szklarczyk, D. et al. STRING v11: protein–protein association networks with increased coverage, supporting functional discovery in genome-wide experimental datasets. Nucleic Acids Res. 47, D607–D613 (2019).

21. Han, H. et al. TRRUST v2: an expanded reference database of human and mouse transcriptional regulatory interactions. Nucleic Acids Res. 46, D380–D386 (2018).

22. Shannon, P. et al. Cytoscape: a software environment for integrated models of biomolecular interaction networks. Genome Res. 13, 2498–2504 (2003).

23. Bhat-Nakshatri, P. et al. A single-cell atlas of the healthy breast tissues reveals clinically relevant clusters of breast epithelial cells. Cell Rep. Med. 2, 100219 (2021).

24. Nguyen, Q. H. et al. Profiling human breast epithelial cells using single cell RNA sequencing identifies cell diversity. Nat. Commun. 9, (2018).

25. Pal, B. et al. A single-cell RNA expression atlas of normal, preneoplastic and tumorigenic states in the human breast. EMBO J. 40, e107333 (2021).

26. Karaayvaz, M. et al. Unravelling subclonal heterogeneity and aggressive disease states in TNBC through single-cell RNA-seq. Nat. Commun. 9, (2018).

27. Costa, A. et al. Fibroblast Heterogeneity and Immunosuppressive Environment in Human Breast Cancer. Cancer Cell 33, 463–479.e10 (2018).

28. Han, C., Liu, T. & Yin, R. Biomarkers for cancer-associated fibroblasts. Biomark. Res. 8, (2020).

29. Zheng, S. et al. Landscape of cancer-associated fibroblasts identifies the secreted biglycan as a protumor and immunosuppressive factor in triple-negative breast cancer. OncoImmunology 11, (2022).

30. Wu, S. Z. et al. A single-cell and spatially resolved atlas of human breast cancers. Nat. Genet. 53, 1334–1347 (2021).

31. Mukund, K. et al. Immune Response in Severe and Non-Severe Coronavirus Disease 2019 (COVID-19) Infection: A Mechanistic Landscape. Front. Immunol. 12, 738073 (2021).

32. Xiong, D., Wang, Y. & You, M. A gene expression signature of TREM2hi macrophages and γδ T cells predicts immunotherapy response. Nat. Commun. 11, (2020).

33. Lin, Y., Xu, J. & Lan, H. Tumor-associated macrophages in tumor metastasis: biological roles and clinical therapeutic applications. J. Hematol. Oncol.J Hematol Oncol 12, (2019).

34. Aran, D., Hu, Z. & Butte, A. J. xCell: digitally portraying the tissue cellular heterogeneity landscape. Genome Biol. 18, (2017).

35. Racle, J. & Gfeller, D. EPIC: A Tool to Estimate the Proportions of Different Cell Types from Bulk Gene Expression Data. Bioinforma. Cancer Immunother. 2120, 233–248 (2020).

36. Luca, B. A. et al. Atlas of clinically distinct cell states and ecosystems across human solid tumors. Cell 184, 5482–5496.e28 (2021).

37. Steen, C. B. et al. The landscape of tumor cell states and ecosystems in diffuse large B cell lymphoma. Cancer Cell 39, 1422–1437.e10 (2021).

38. Lujano Olazaba, O., Farrow, J. & Monkkonen, T. Fibroblast heterogeneity and functions: insights from single-cell sequencing in wound healing, breast cancer, ovarian cancer and melanoma. Front. Genet. 15, 1304853 (2024).

39. Kloosterman, D. J. & Akkari, L. Macrophages at the interface of the co-evolving cancer ecosystem. Cell 186, 1627–1651 (2023).

40. Hammerl, D. et al. Spatial immunophenotypes predict response to anti-PD1 treatment and capture distinct paths of T cell evasion in triple negative breast cancer. Nat. Commun. 12, (2021).

41. Jiang, W. et al. Exhausted CD8+T Cells in the Tumor Immune Microenvironment: New Pathways to Therapy. Front. Immunol. 11, (2021).

42. Tian Wei-hong et al. Hierarchical transcriptional network governing heterogeneous T cell exhaustion and its implications for immune checkpoint blockade. Front. Immunol. 14, (2023).

43. Mariathasan, S. et al. TGFβ attenuates tumour response to PD-L1 blockade by contributing to exclusion of T cells. Nature 554, 544–548 (2018).

44. Frigo, D. E., Bondesson, M. & Williams, C. Nuclear receptors: from molecular mechanisms to therapeutics. Essays Biochem. 65, 847–856 (2021).

45. Borin, T., Angara, K., Rashid, M., Achyut, B. & Arbab, A. Arachidonic Acid Metabolite as a Novel Therapeutic Target in Breast Cancer Metastasis. Int. J. Mol. Sci. 18, 2661 (2017).

46. Borin, T. F. et al. HET0016 decreases lung metastasis from breast cancer in immune-competent mouse model. PLOS ONE 12, e0178830 (2017).

47. Elsayed, A. A. R., Al-Marsoummi, S., Vomhof-Dekrey, E. E. & Basson, M. D. Slfn12 over-expression sensitizes triple negative breast cancer cells to chemotherapy drugs and radiotherapy. Cancer Genomics Proteomics 19, 328–338 (2022).

48. Singhal, S. K. et al. Schlafen 12 slows tnbc tumor growth, induces luminal markers, and predicts favorable survival. Cancers 15, 402 (2023).

49. Ashour, N. et al. Epigenetic regulation of gfi1 in endocrine-related cancers: a role regulating tumor growth. Int. J. Mol. Sci. 21, 4687 (2020).

50. Sun, L. et al. Advances in understanding the roles of cd244 (Slamf4) in immune regulation and associated diseases. Front. Immunol. 12, (2021).

51. Wherry, E. J. & Kurachi, M. Molecular and cellular insights into T cell exhaustion. Nat. Rev. Immunol. 15, 486–499 (2015).

52. Cheng, J., Ding, X., Xu, S., Zhu, B. & Jia, Q. Gene expression profiling identified TP53MutPIK3CAWild as a potential biomarker for patients with triple-negative breast cancer treated with immune checkpoint inhibitors. Oncol. Lett. 19, 2817–2824 (2020).

53. Liverani, C. et al. Lineage-specific mechanisms and drivers of breast cancer chemoresistance revealed by 3D biomimetic culture. Mol. Oncol. 16, 921–939 (2021).

54. Arcucci, A., Ruocco, M. R., Granato, G., Sacco, A. M. & Montagnani, S. Cancer: an oxidative crosstalk between solid tumor cells and cancer associated fibroblasts. BioMed Res. Int. 2016, 1–7 (2016).

55. Mollah, F. & Varamini, P. Overcoming therapy resistance and relapse in tnbc: emerging technologies to target breast cancer-associated fibroblasts. Biomedicines 9, 1921 (2021).

56. Tan, Q. et al. Breast cancer cells interact with tumor-derived extracellular matrix in a molecular subtype-specific manner. Biomater. Adv. 146, 213301 (2023).

57. Yeh, Y.-H., Hsiao, H.-F., Yeh, Y.-C., Chen, T.-W. & Li, T.-K. Inflammatory interferon activates HIF-1α-mediated epithelial-to-mesenchymal transition via PI3K/AKT/mTOR pathway. J. Exp. Clin. Cancer Res. 37, (2018).

58. Liu, Q. et al. Targeting hypoxia-inducible factor-1alpha: A new strategy for triple-negative breast cancer therapy. Biomed. Pharmacother. 156, 113861 (2022).

59. Zhao, L., Zhou, S. & Gustafsson, J.-Å. Nuclear receptors: recent drug discovery for cancer therapies. Endocr. Rev. (2019) doi:10.1210/er.2018-00222.

60. Yousefi, H., Fong, J. & Alahari, S. K. Nr4a family genes: a review of comprehensive prognostic and gene expression profile analysis in breast cancer. Front. Oncol. 12, 777824 (2022).

61. Muscat, G. E. O. et al. Research resource: nuclear receptors as transcriptome: discriminant and prognostic value in breast cancer. Mol. Endocrinol. 27, 350–365 (2013).

62. Zhou, F. et al. Nuclear receptor NR4A1 promotes breast cancer invasion and metastasis by activating TGF-β signalling. Nat. Commun. 5, 3388 (2014).

63. Wang, D. & DuBois, R. N. The Role of Prostaglandin E 2 in Tumor-Associated Immunosuppression. Trends Mol. Med. 22, 1–3 (2016).

64. Hossain, F. et al. Sulindac sulfide as a non-immune suppressive γ-secretase modulator to target triple-negative breast cancer. Front. Immunol. 14, 1244159 (2023).

