## Supplementary Information for "A novel subtyping method for TNBC with implications for prognosis and therapy"

**Supplementary Table 1:** List of all of the cell type-markers.

| Basal | NDUFA4L2 | CAFs | ACTA2 | B cell | MS4A1 |
| --- | --- | --- | --- | --- | --- |
|  | CD36 |  | PDGFRA |  | CD79A |
|  | VIM |  | PDGFRB |  | CD79B |
|  | ACKR1 |  | PDNPN |  | BLNK |
|  | IL8 |  | THY1 |  | IGHD |
|  | MECOM |  | COL1A1 |  | IGHM |
|  | CLDN5 |  | CAV1 |  | IGHA1 |
|  | TCF4 |  | ITGB1 |  | IGHA2 |
|  | ZEB1 |  | S100A4 |  | IGKV |
|  | Notch4 |  | AIFM2 |  | IGLC3 |
|  | DKK3 |  | BSG | Plasmablast | CD27 |
|  | KRT14 |  | CD74 |  | CD38 |
|  | ITGA6 |  | C5AR2 | Follicular B cell | MEF2B |
|  | KRT5 |  | CD70 |  | BCL6 |
|  | TP63 |  | ALDH1A1 |  | RGS13 |
|  | KRT17 |  | LEPR |  | NEIL1 |
|  | MME |  | TAGLN | Monocyte | CD14 |
|  | SOX2 |  | FAP |  | FCGR3A |
|  | NANOG |  | DAN |  | HLA-DRA |
|  | KLF4 | Fibroblast | MMP2 |  | TNFRSF1B |
|  | IRX4 |  | APOD |  | CD33 |
|  | MEF2C |  | CLMP |  | SELL |
|  | SLUG |  | MMP3 | Macrophage | CSF1R |
|  | ERG2 |  | DPT |  | CD68 |
|  | TWIST2 |  | IGFBP6 |  | CD163 |
|  | TBX2 |  | DCN | M1 Macrophage | IL12A |
| Pan Luminal | KRT8 |  | LUM |  | TNF |
|  | KRT18 |  | SNAI1 |  | IL6 |
| Luminal Progenitor | SFRP1 |  | ZEB2 |  | IL1B |
|  | CXXC5 |  | POSTN |  | IL23A |
|  | SLP1 |  | COL3A1 |  | NFKB1 |
|  | ANXA1 |  | SPARC |  | STAT1 |
|  | RARRES1 |  | COL5A2 |  | IRF5 |
|  | KLK5 |  | COL14A1 | M2 Macrophage | ARG1 |
|  | KRT15 |  | COL10A1 |  | ARG2 |
|  | SCGB2A1 |  | WISP2 |  | STAT6 |
|  | CALML5 |  | CILP |  | MMP14 |
|  | GLYATL2 |  | SFRP4 |  | MMP9 |
|  | TOP2A |  | COMP |  | CD276 |
|  | NUSAP1 |  | NFAT |  | FN1 |
|  | UBE2C |  | HIF1A |  | MRC1 |
|  | TPX2 |  | EPAS1 |  | CCL13 |
|  | SPC25 |  | FLI1 |  | CCL18 |
|  | MKI67 |  | FOS |  | LYVE1 |
|  | CDK1 |  | JUN |  | PDCD1LG2 |
|  | CENPF |  | EGR |  | TGFB2 |
|  | CCNA2 |  | ATF3 |  | IL10 |
|  | MEG3 |  | GADD45 |  | CCL2 |
|  | IGF1 |  | HSP1A1 | TAM | TREM1 |
|  | PTGDS |  | DNAJB |  | TREM2 |
|  | KIT | Endothelial | VWF |  | CD1D |
|  | ALDH1A3 |  | KDR |  | ITGAM |
|  | EHF |  | VWF |  | VCAM1 |
|  | ELF5 |  | KDR |  | VTCN1 |
|  | CYP24A1 |  | CDH5 |  | STAT3 |
|  | LBP |  | VCAM1 |  | HLA-DRB1 |
|  | SOX10 |  | VEGF |  | MSR1 |
| Mature Luminal | ESR1 |  | KLF4 | NK cell | CD7 |
|  | TNFSF11 |  | NOS3 |  | PRF1 |
|  | PGR | Immature pericytes | CD44 |  | NKG7 |
|  | FOXA1 |  | CSPG4 |  | GZMA |
|  | TBX3 |  | RGS5 |  | GZMB |
|  | PDK4 | γδ T cell | CD5 |  | GZMK |
|  | XBP1 |  | TRDC |  | ACTB |
|  | STC2 |  | TRGV9 |  | ARPC3 |
|  | RUNX1 | CD4 T cell | CD4 |  | CFL1 |
|  | BATF |  | CCR7 |  | CST7 |
|  | ANKRD30A |  | LEF1 |  | KIR2DL1 |
|  | PIP |  | TCF7 |  | KIR2DL3 |
|  | MUCL1 |  | CTLA4 |  | KIR3DL1 |
|  | TAT |  | FOXP3 |  | KIR3DL2 |
|  | TSPAN8 |  | GATA3 |  | KLRC2 |
|  | PRLR |  | IL2R |  | IL32 |
|  | NFE2L2 |  | ICOS |  | GZMH |
|  | CHD1 | CD8 T cell | PTPRC |  | XCL1 |
|  | MYC |  | CD2 |  | KLRC1 |
|  | LMO4 |  | CD3D |  | KLRB1 |
|  | MYB |  | CD3E |  | SREG |
|  | PR |  | CD3G |  | FCER1G |
|  | CITED1 |  | CD8A | Neutrophil | CD11A |
| Myoepithelial | TGLN |  | CD8B |  | CD11B |
|  | MYLK |  | CCL5 |  | ITGAL |
|  | ACTG2 |  | IFNG |  | CD55 |
|  | ACTN1 |  | CCL3 |  | CD16 |
|  | CALD1 |  | CCL4 |  | CD10 |
|  | MYL9 |  | PDCD1 |  |  |
|  | RP63 |  | LAG3 |  |  |
|  | PPARG |  | TIGIT |  |  |

**Supplementary Figure 1: Distribution of self-reported race (SRR) and genetic ancestry across TNBC subtypes.**

Features a pie chart showing the proportion of SRR in each TNBC subtype (top panels). Features a pie chart showing the proportion of AA and EA who have >60% Central/West African and Western/Central European ancestry, respectively, in each subtype (bottom panels).

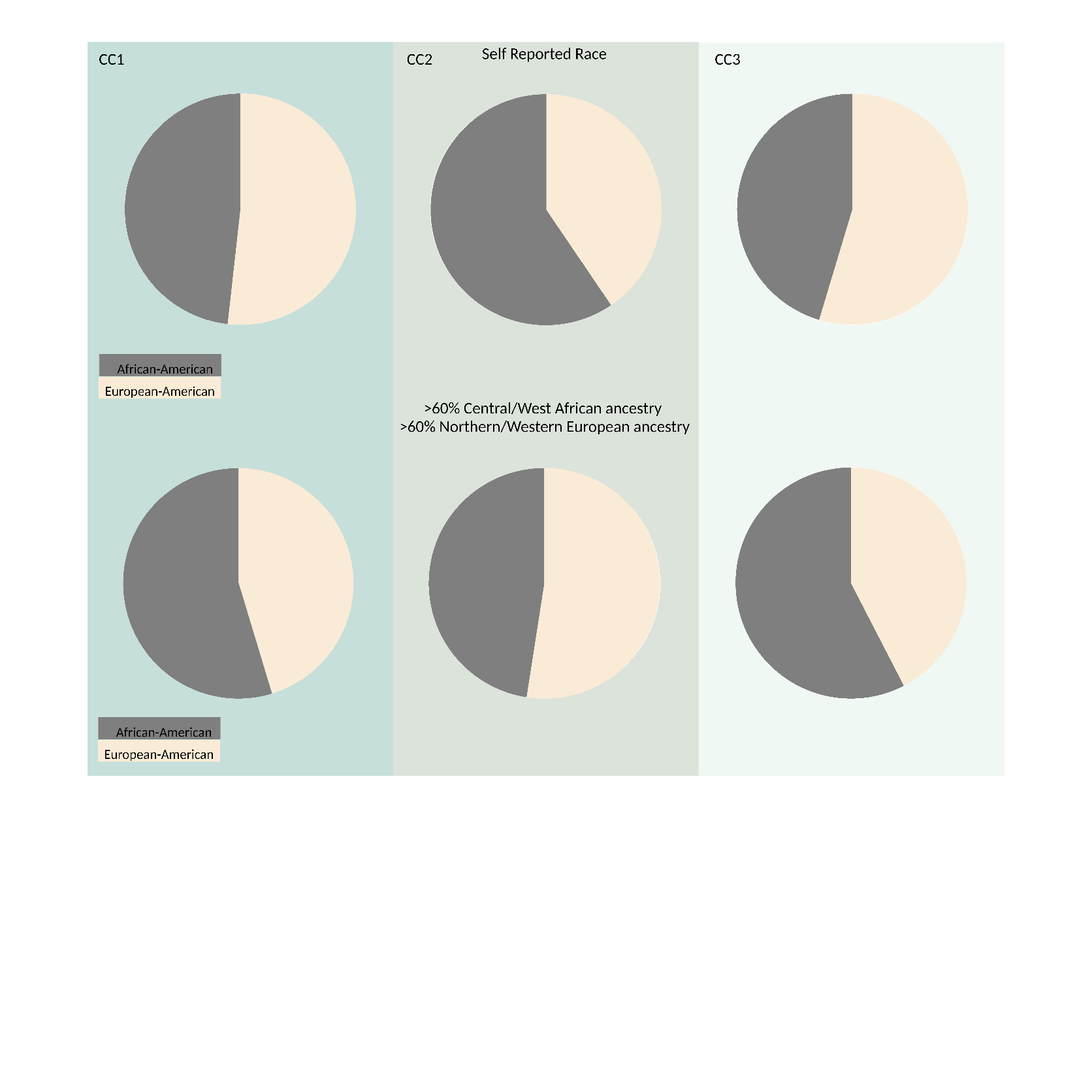

**Supplementary Figure 2: Cellular composition analysis across TNBC subtypes and groups.**

**A.** A heatmap is displaying the relative enrichment score for each of the xCELL 64 cell types across the samples in each subtype (ConsensusCluster) and group (HierarchicalCluster) in the LTR TNBCs. **B.** A barplot shows the proportion of different EPIC cell types in each sample. **C.** This table encapsulates the results from xCELL, EPIC, and EcoTyper for each of the subtypes.

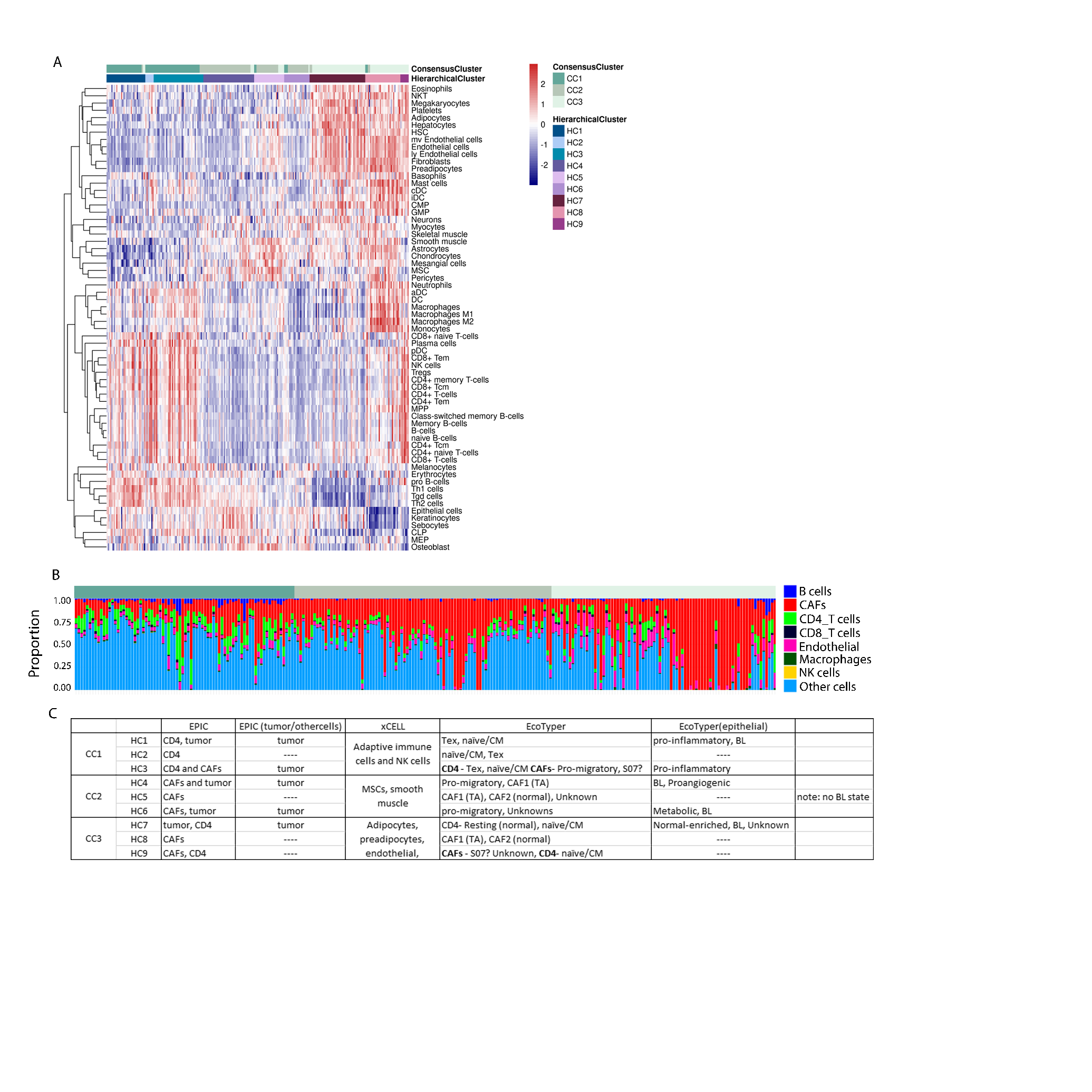

**Supplementary Figure 3:** Enrichment of GO ontology gene sets, for genes differentially expressed in each subtype compared to the others.

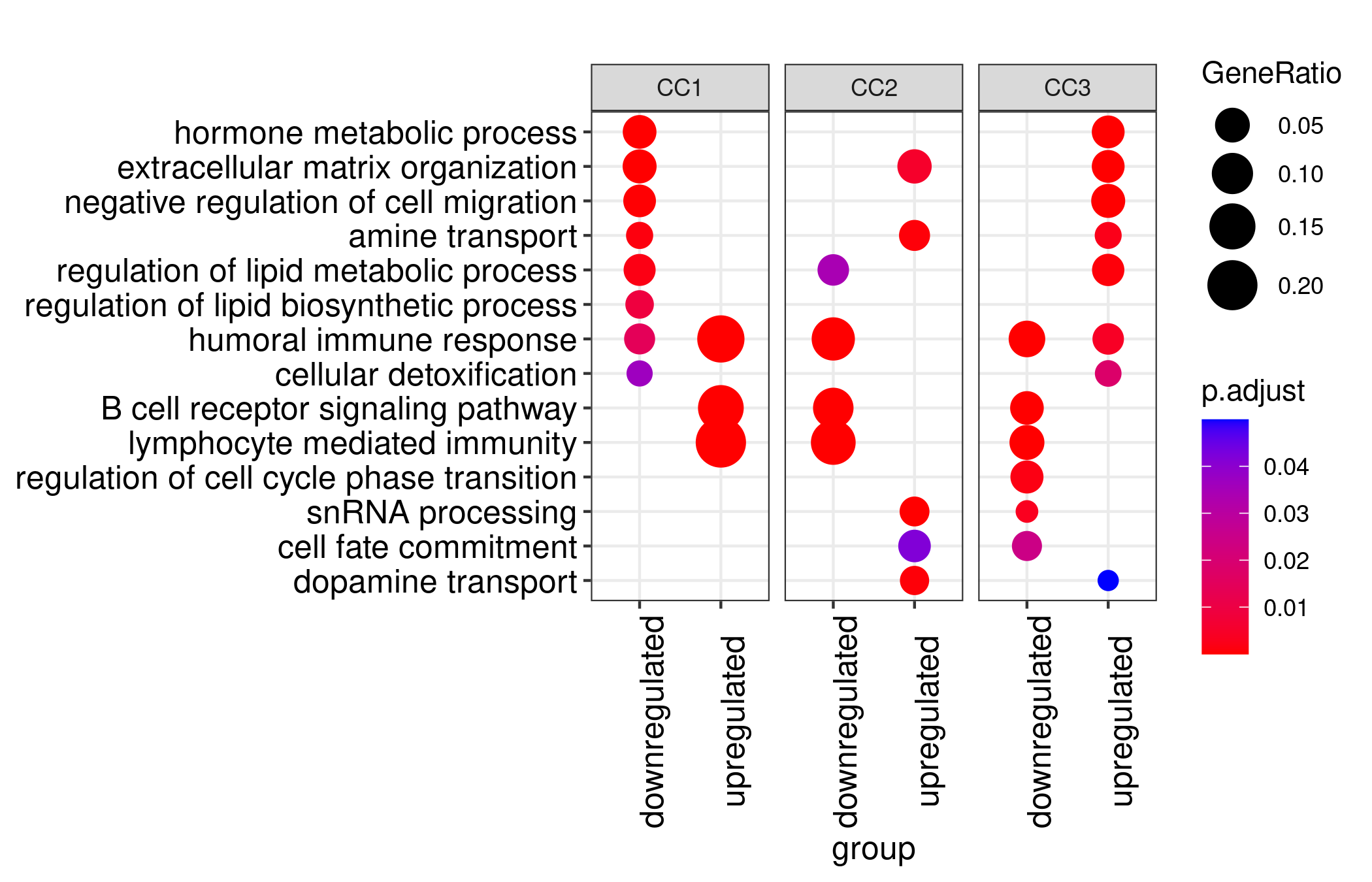

**Supplementary Figure 4: Ancestry-inferred subtype functional differences cnetplot.**

**A.** A CC1 specific network enriched in AA samples, highlighting immunoglobulin production and antigen recognition. **B.** A CC1 specific network in EA samples, highlighting embryonic development and stemness. **C.** A CC2 specific network in AA samples, highlighting humoral immunity. **D.** A CC3 specific network in AA samples, highlighting immunoglobulin production and antigen presentation via MHC-II. and **E.** A CC3 specific network in EA samples, highlighting epithelial cell processes and metastasis.

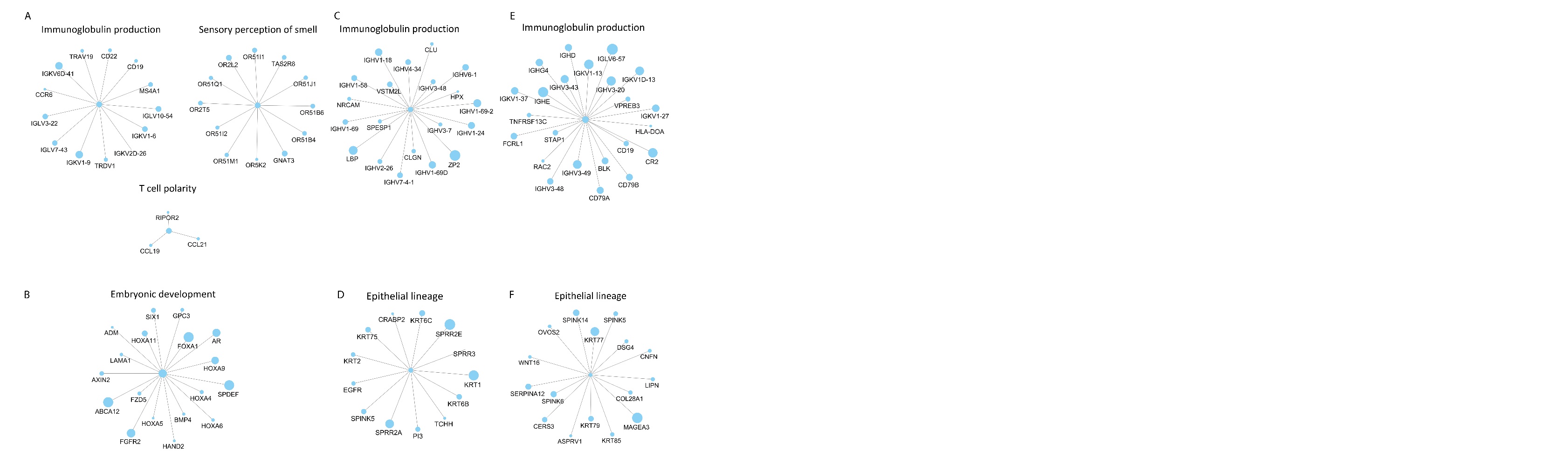

**Supplementary Figure 5: Nuclear receptor expression and CC3-specific network analysis in LTR TNBCs.**

**A.** A heatmap displays the relative gene expression of 48 members of the human nuclear receptors in each subtype (ConsensusCluster) and group (HierarchicalCluster) in the LTR TNBCs. **B.** A CC3-specific network captures nuclear receptors and its neighboring DEGs. Nodes are sized and colored based on the log2FC value between subtypes: dark blue (log2FC of -2) to white (0) to dark red (2). The edges represent protein-protein interactions, with arrow indicating transcription to target relationships. The teal boarder color indicates nuclear receptor markers.

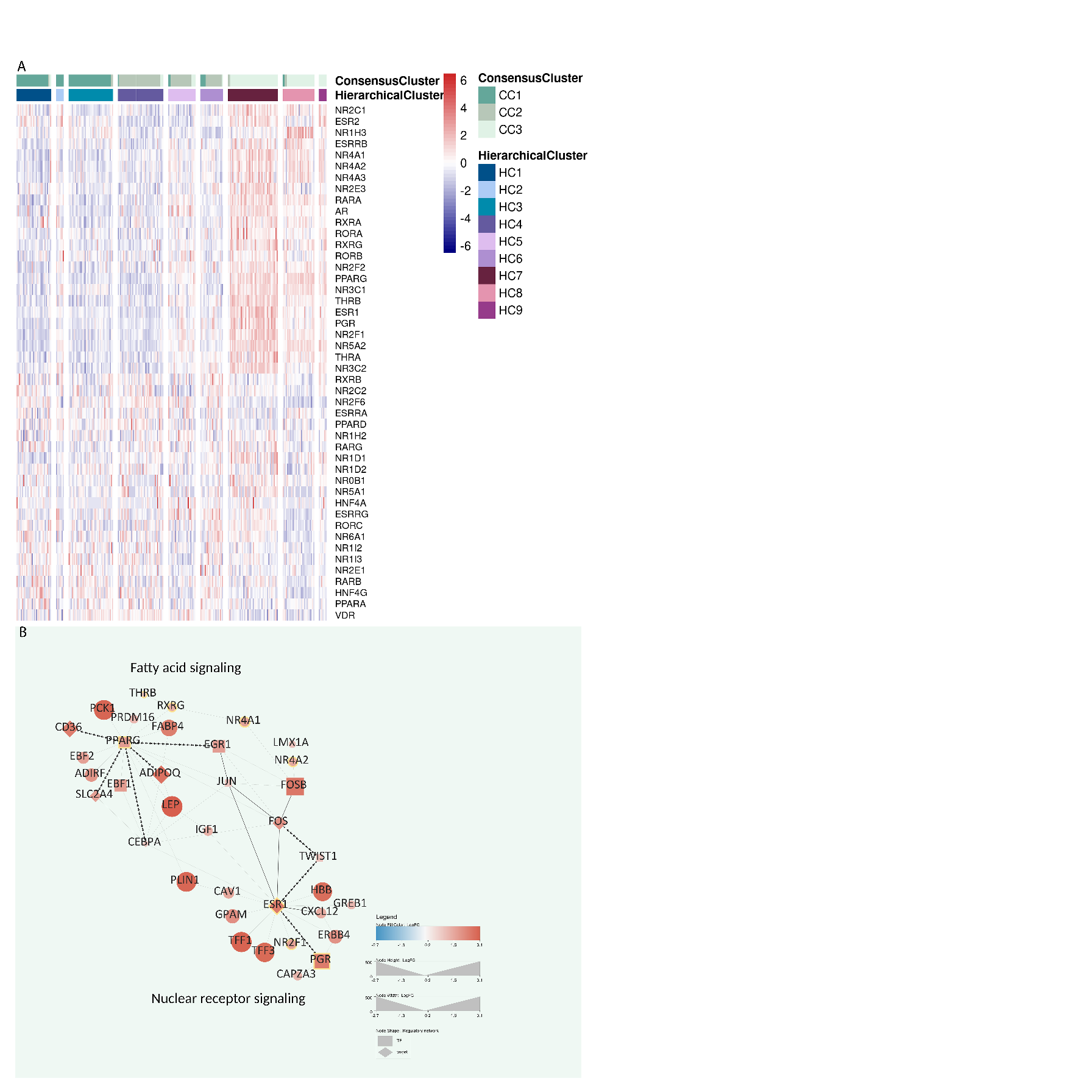

**Supplementary Figure 6: Histopathology and Immunohistochemistry in a representative case of TNBC.**

Three areas of the tumor are shown. Areas of solid tumors are characterized by nests of poorly differentiated epithelial cells, with hyperchromatic nuclei; in these areas, expression of Estrogen Receptor is completely negative (Left Panels). Towards the periphery of the tumor, a few normal breast glands are trapped by neoplastic cells, showing significant lymphocytic infiltration; in these areas, the tumor remains negative and the trapped glands show moderate nuclear expression of ER (middle panels). On the edge, the tumor cells are separated from normal glands, which show robust nuclear labeling of ER (right panels). All panels' original magnification is 200x.

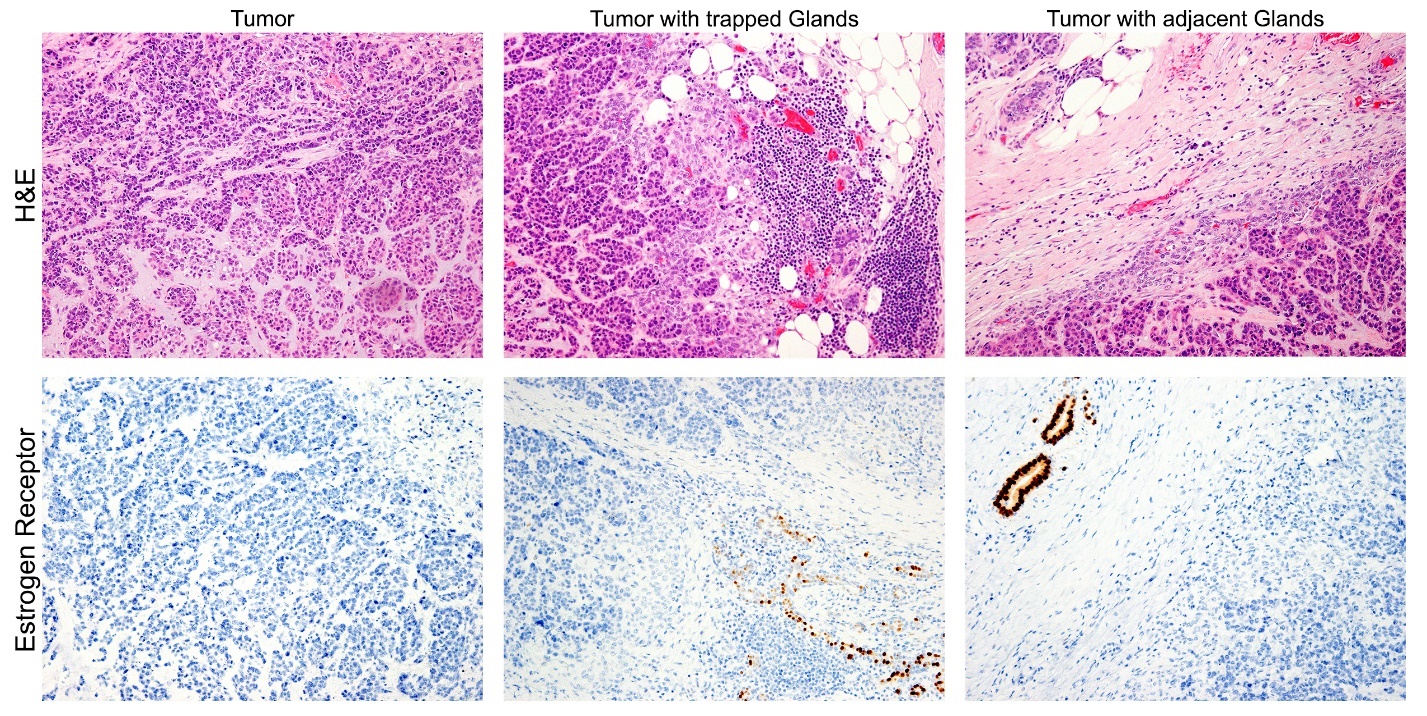

**Supplementary Figure 7:** A heatmap displays the relative gene expression of hypoxia-specific markers in each subtype (ConsensusCluster) and group (HierarchicalCluster) in the LTR TNBCs.

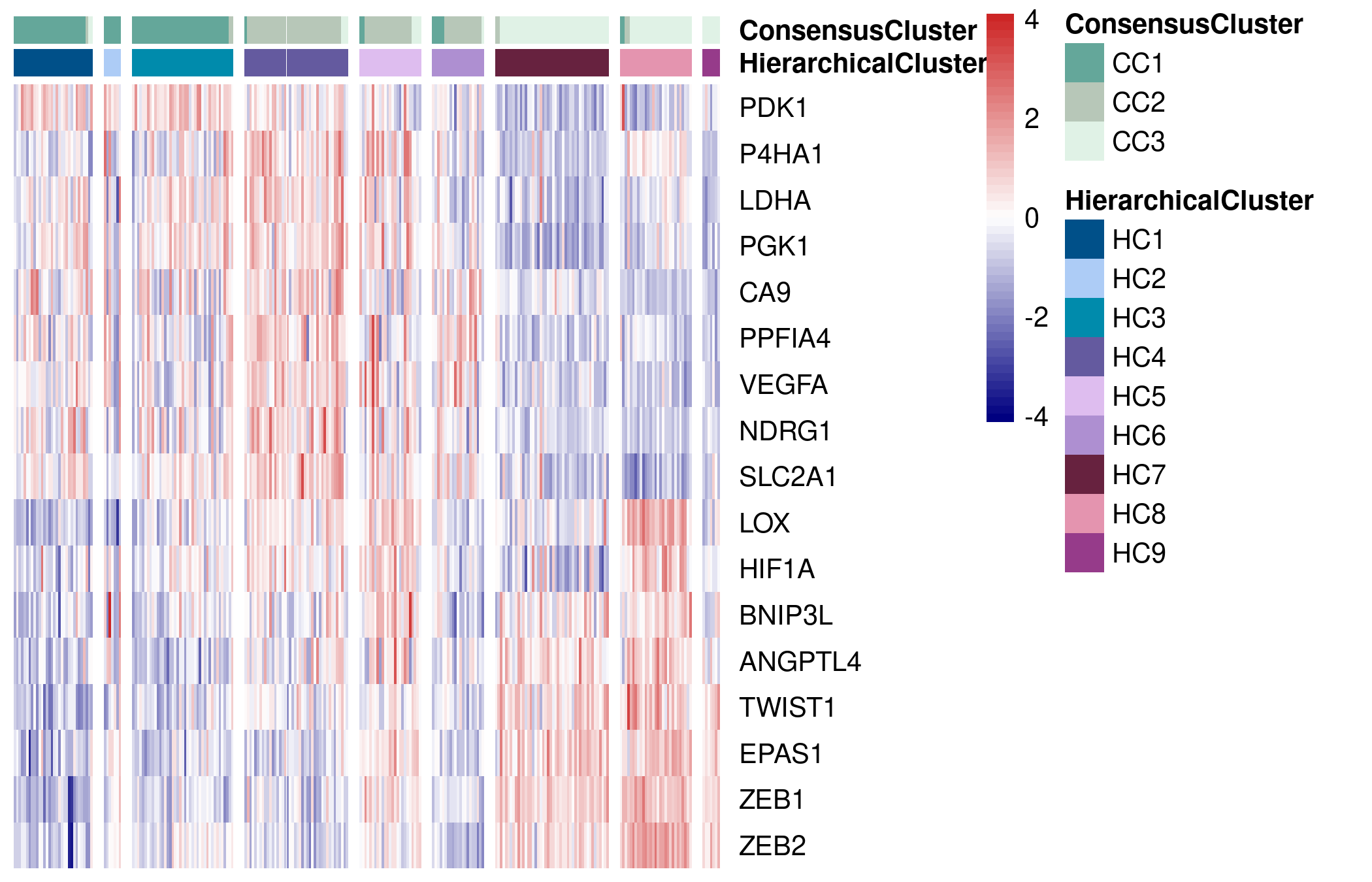

**Supplementary Figure 8:** Boxplots show the scaled expression of PTGES, PTGS1, and PTGS2 across each TNBC subtypes.

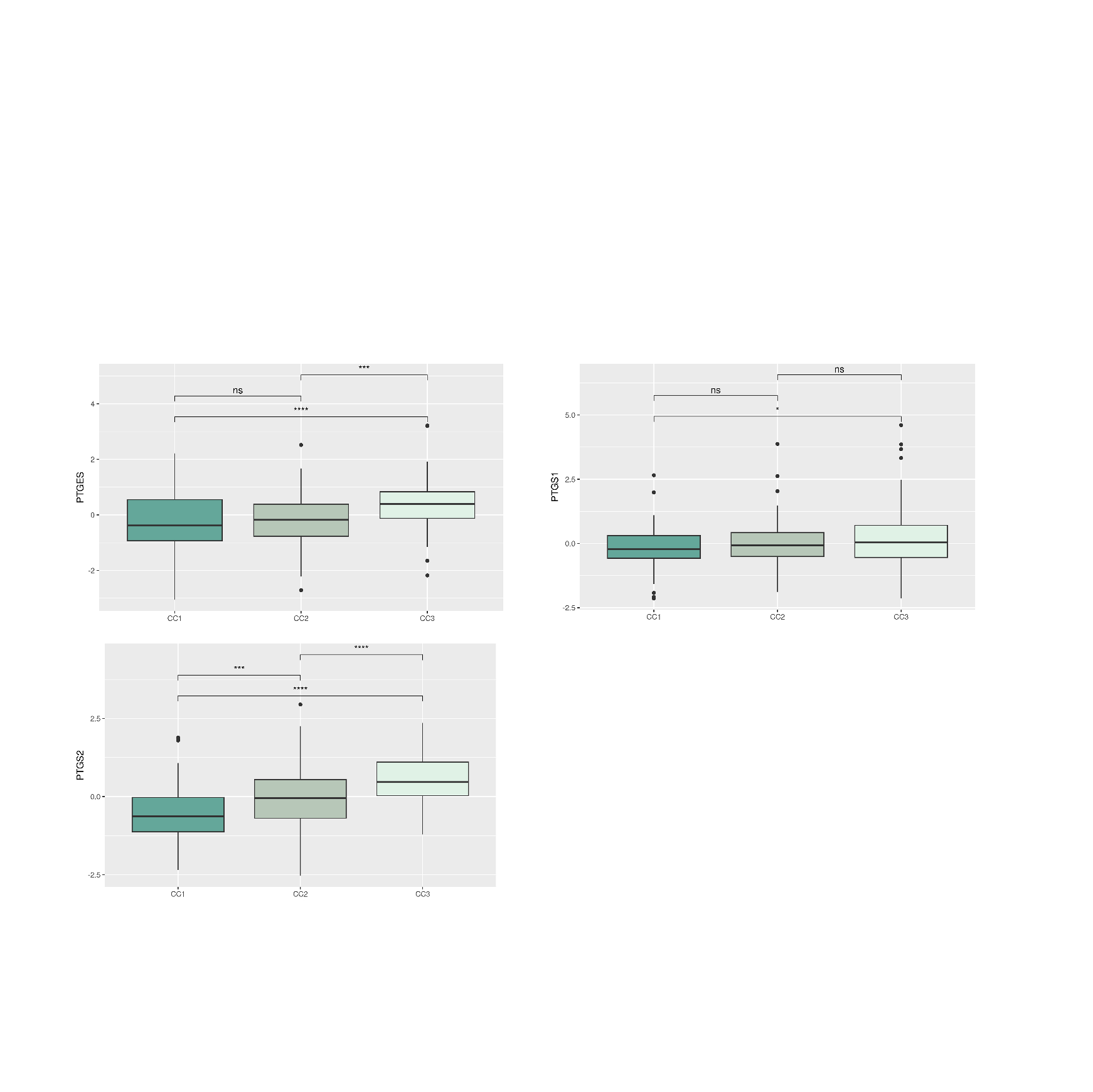
